# Heimdall SNAREs reveal an ancestral architecture of eukaryotic SNARE complexes

**DOI:** 10.64898/2026.10.19.760048

**Authors:** Iman Bentahar, Deepak Yadav, Aysima Hacisuleyman, Dirk Fasshauer

## Abstract

SNARE proteins are central to eukaryotic cell biology, where they drive membrane fusion by assembling into four-helix bundles that bring opposing membranes into close proximity. Their ubiquitous role in intracellular membrane trafficking raises fundamental questions about their evolutionary origin and the emergence of eukaryotic cellular complexity. Here, we characterize SNARE-like proteins from Heimdallarchaeota, a lineage within the Asgard archaea that is closely related to eukaryotes. We show that a Heimdall SNARE forms a stable homotetrameric four-helix bundle in solution and determine its structure by X-ray crystallography. Strikingly, the Heimdall complex closely resembles eukaryotic SNARE complexes in its overall architecture and core organization, despite being composed of four identical subunits. Molecular dynamics simulations further reveal conserved physical features of the bundle alongside distinctive properties of its central 0-layer. These findings demonstrate that the characteristic four-helix architecture of SNAREs predates the emergence of eukaryotic cells and provide structural insight into the evolution of the membrane fusion machinery.

## Introduction

The origin of the complex eukaryotic cell remains one of the central puzzles in evolutionary biology ^1–4^; ^5^. Compared to prokaryotes, eukaryotic cells are considerably larger and are compartmentalized by membrane-bound organelles such as mitochondria, the nucleus, and a sophisticated endomembrane system. While the mitochondrial endosymbiont was long ago recognized as a former bacterium ^6,7^, the evolutionary origin of the endomembrane system—which includes the nuclear envelope, endoplasmic reticulum (ER), Golgi apparatus, lysosomes, vacuoles, and plasma membrane—is still unclear. Intracellular transport within this system is primarily mediated by vesicles that move along dynamic cytoskeletal tracks such as microtubules and actin filaments, powered by motor proteins ^8–10^. The evolution of a dynamic cytoskeleton was itself a key innovation in early eukaryotes, enabling not only intracellular trafficking but also cell motility.

The discovery of Asgard archaea through metagenomic studies of marine sediments ^11^ has provided new insights into the transition from prokaryotic to eukaryotic cell architecture. These archaea, named after Norse gods (e.g., Loki, Thor, Odin), are notable for possessing genes previously thought to be eukaryote-specific, including components related to membrane trafficking and cytoskeletal regulation ^11–14^. The presence of these so-called eukaryotic signature proteins (ESPs) in Asgard archaea suggests a shared ancestry with eukaryotes. Structural and biochemical studies have already provided insight into the cytoskeleton of Asgard archaea, including tubulin ^15,16^ and actin homologs and their regulatory partners, such as profilin and gelsolin^17–20^. Importantly, cultivation-based studies have begun to link this molecular repertoire to cellular phenotypes: cultivated Lokiarchaea exhibit elaborate protrusions and a highly flexible cell architecture ^21,22^, consistent with a dynamic cytoskeletal system. Moreover, two recently cultivated Heimdallarchaeota species display intracellular vesicular structures ^23,24^, raising the possibility that a simple, archaeal precursor of eukaryotic membrane trafficking systems may have already been present in this lineage.

Asgard proteins implicated in membrane remodelling and trafficking—such as the ESCRT-III complex ^25–27^, several types of small GTPases of the Ras superfamily, among them Sar1 ^28^; ^29^, the inner coat protein Sec23 ^12^; ^30^, and Soluble N-ethylmaleimide-sensitive factor Attachment protein REceptor (SNARE) proteins ^31^—have clear homology to their eukaryotic counterparts. In eukaryotes, different sets of SNARE proteins catalyze the fusion of different transport vesicles with their respective target compartments. Eukaryotic SNARE complexes form stable four-helix bundles composed of four distinct SNARE motifs (Qa, Qb, Qc, and R), which zipper together to drive membrane fusion ^32–34^. The mechanism was first elucidated through studies of neurotransmitter release in neurons (reviewed in ^35–37^; ^38^).

In the present study, we extend this work by solving the crystal structure of a SNARE-like protein from Heimdallarchaea. The protein forms a homotetrameric four-helix bundle that is highly similar to eukaryotic SNARE complexes in both structure and stability. These findings support the hypothesis that the ancestral eukaryotic SNARE complex evolved from a homomeric precursor encoded in an archaeal genome.

## Results

### Stability and oligomeric state of the Heimdall SNARE complex

In a previous study, we showed that the purified SNARE-like domains from *Heimdallarchaeota*, referred to as HPS1 and HPS2, adopt an α-helical conformation as assessed by circular dichroism (CD) spectroscopy ^31^, consistent with a coiled-coil structure. To assess the stability of this structural arrangement, we monitored thermal unfolding by gradually heating the purified proteins. Both HPS1 and HPS2 exhibited remarkable thermal stability, with unfolding transitions occurring only at temperatures exceeding 90 °C (Fig. 1A). To further probe their structural stability, we examined the effect of chemical denaturants. Urea treatment had little effect on the CD spectra, indicating that the α-helical conformation was largely preserved. In contrast, both proteins lost their α-helical signature upon exposure to 1 M guanidinium hydrochloride (GdnHCl), indicating a cooperative unfolding transition under these conditions (Fig. S1).

**Figure 1:**
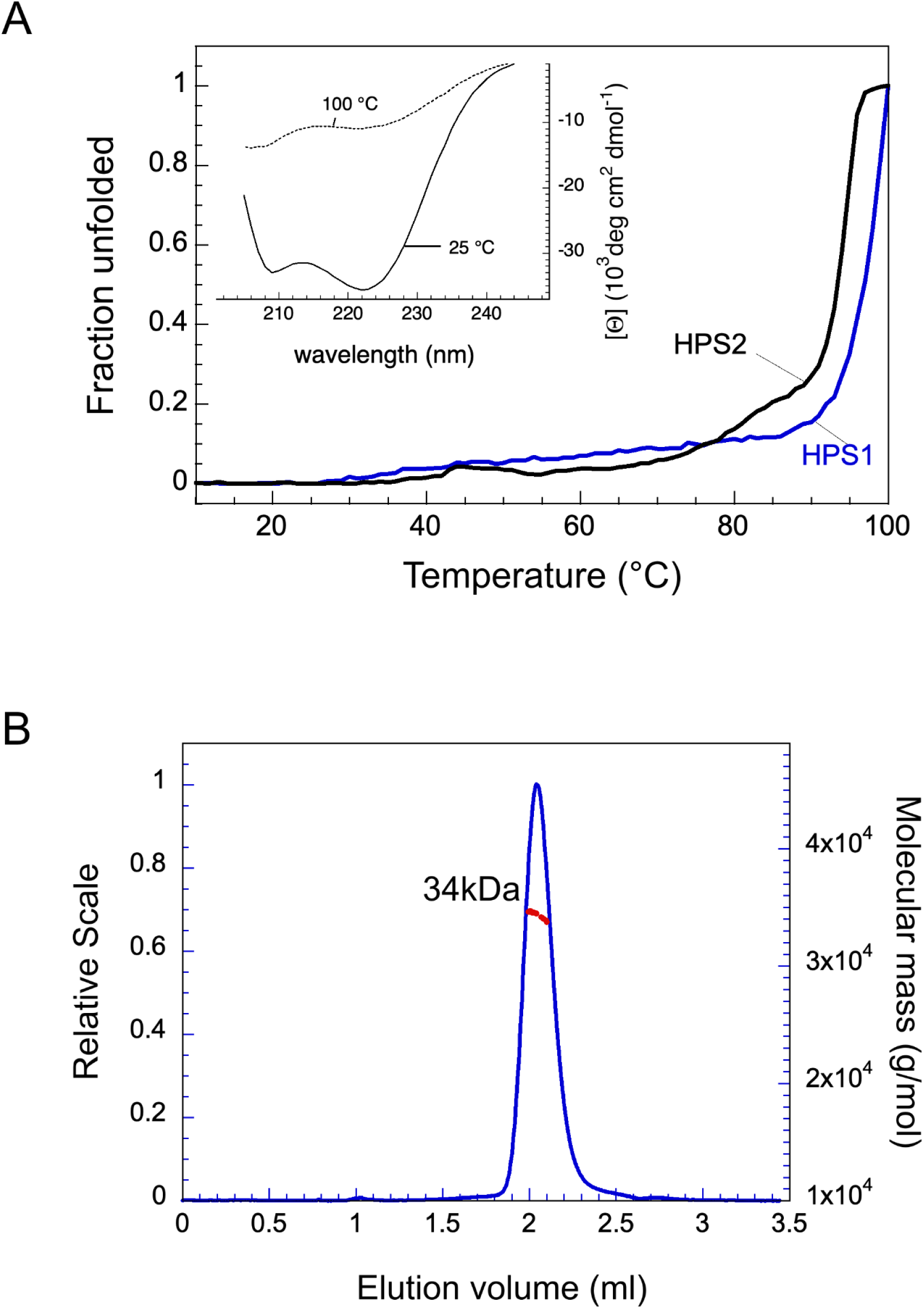
Biophysical characterization of Heimdallarchaeal SNARE-like proteins. **(A)** Thermal unfolding of HPS1 (blue) and HPS2 (black) monitored by circular dichroism at 220 nm from 10–100 °C. HPS2 displays a cooperative transition (Tm ∼95 °C), whereas HPS1 does not fully unfold. Inset: CD spectra before and after thermal denaturation, with the pre-melt spectrum consistent with an α-helical structure. Protein concentrations were 21.5 µM (HPS1) and 20.8 µM (HPS2) in 20 mM Na₃PO₄ (pH 7.4), 150 mM NaCl. **(B)** SEC–MALS analysis of HPS1. A phenylalanine-to-tryptophan substitution (HPS1^F155W) enabled UV detection. The elution profile (blue) shows a single peak at 2.05 mL with a molar mass of ∼34 kDa (red), consistent with a tetramer. The protein was analyzed in 20 mM HEPES (pH 7.4), 150 mM NaCl at a concentration of 4 mg mL^⁻¹^.

The pronounced α-helical character and resistance to denaturation suggested that HPS1 and HPS2 form stable oligomeric assemblies. To determine their oligomeric state in solution, we analyzed both proteins by size-exclusion chromatography (SEC). Both eluted as monodisperse peaks. When calibrated against globular protein standards, HPS1 and HPS2 eluted at apparent molecular masses of ∼30 and ∼70 kDa, respectively (Fig. S2). For HPS1, the apparent mass is consistent with a homotetramer; however, because the protein is expected to adopt an elongated, helical structure, molecular-mass estimates based on globular standards must be interpreted with caution. To obtain a more accurate determination of the oligomeric state, we subjected HPS1 to SEC coupled with multi-angle light scattering (SEC-MALS). This analysis revealed a molecular mass of 34 kDa (Fig. 1B), consistent with a homotetrameric assembly. Taken together, these results suggest that HPS1 forms a stable tetramer in solution, representing its predominant oligomeric state under the conditions tested.

### Crystal structure of the Heimdall SNARE complex reveals a symmetric homotetrameric coiled coil

To further characterize the HPS1 oligomer, we determined the crystal structure of the HPS1 SNARE motif by X-ray crystallography. The crystals belonged to a tetragonal space group with fourfold rotational symmetry, consistent with the homotetrameric assembly observed in solution (Table 1). The structure was determined by molecular replacement using Phaser-MR within the PHENIX suite, with the SNARE motif of human Syntaxin-6 from the early endosomal SNARE complex (PDB ID: 2NPS, chain D) ^39^. Following successful phasing, iterative rounds of model building and refinement yielded a final structure at 2.4 Å resolution.

**Table 1.**
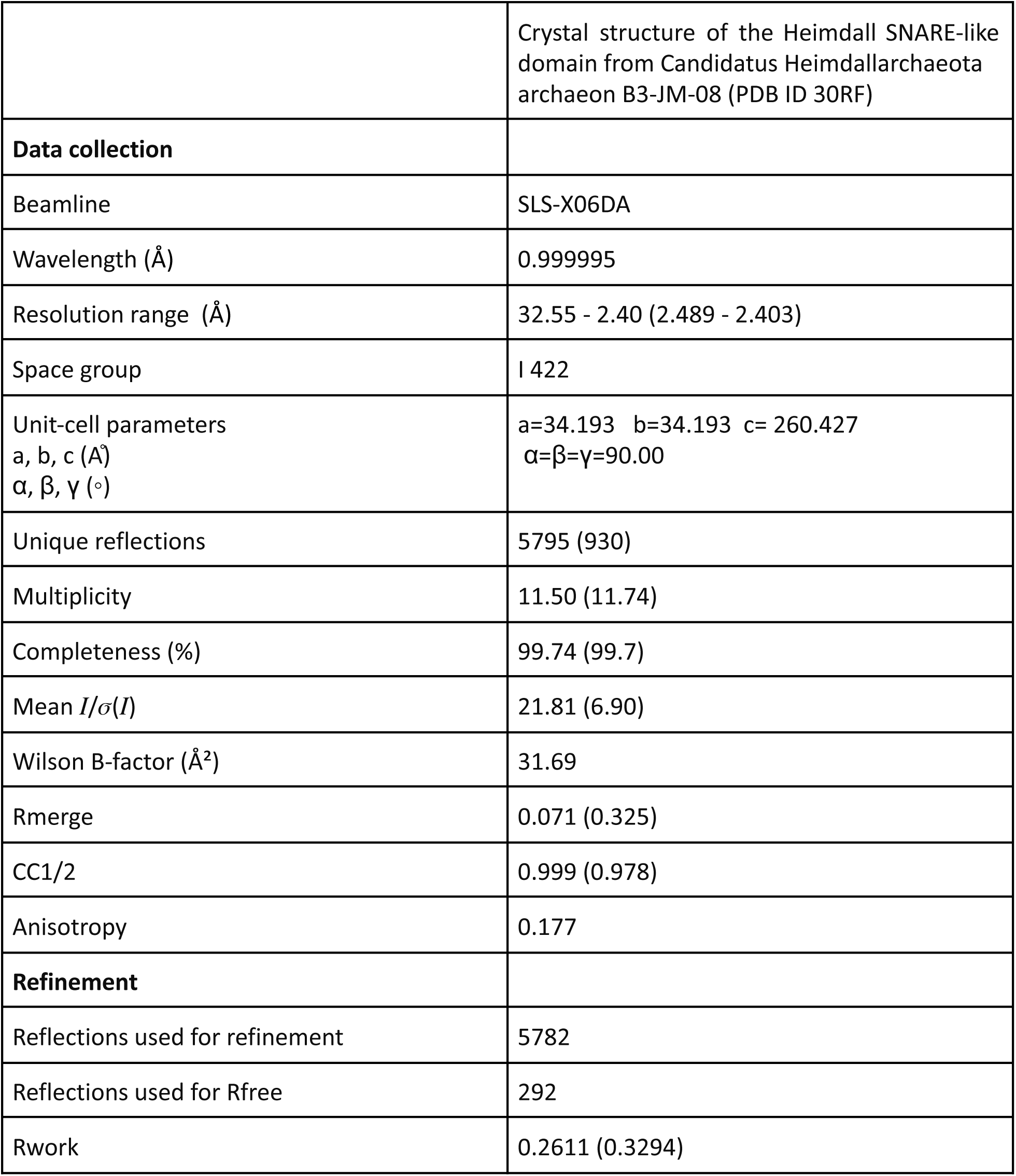
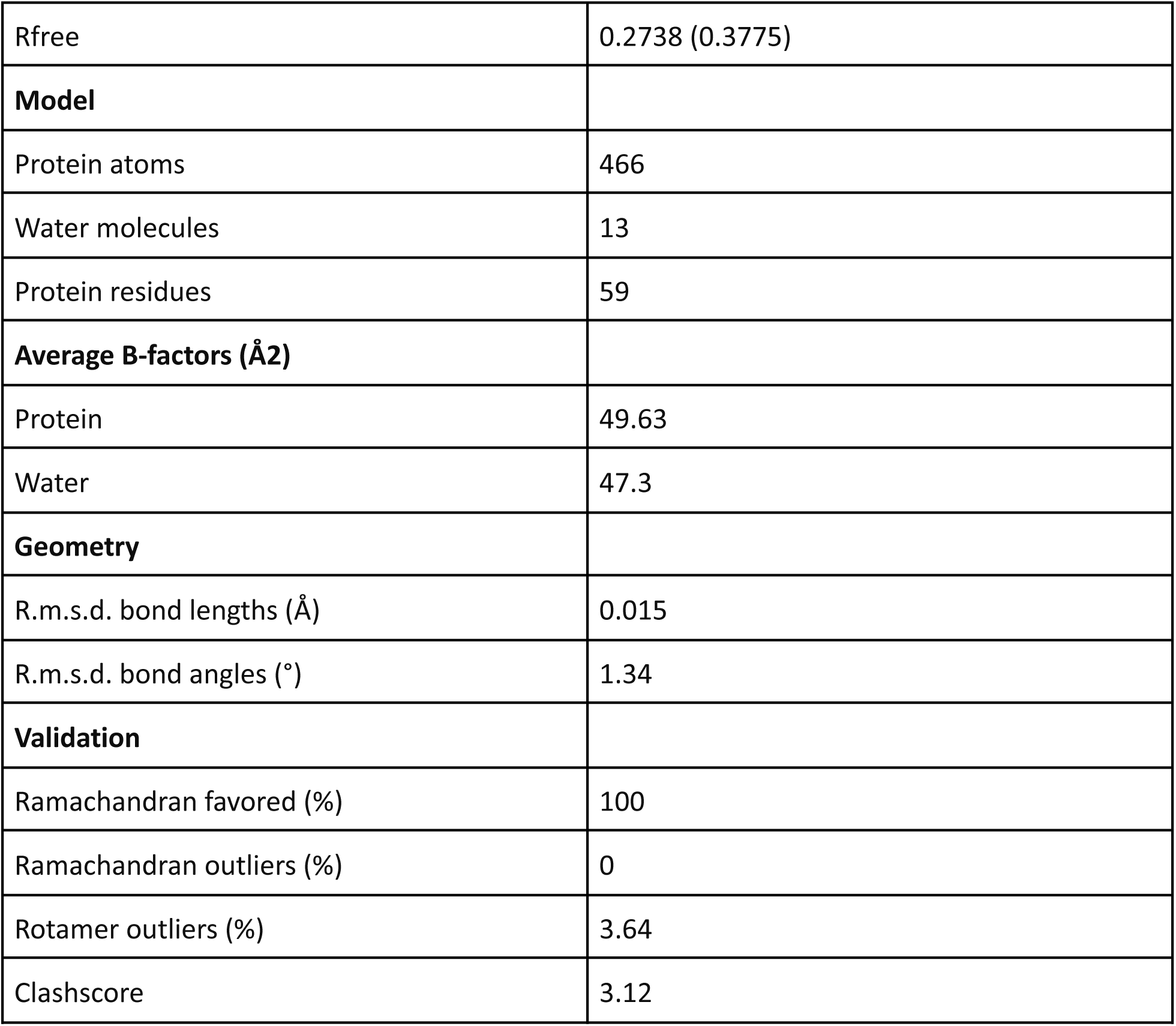
Data collection & refinement statistics. Values for the outer shell are given in parentheses.

|  |  |
| --- | --- |
|  | Crystal structure of the Heimdall SNARE-like domain from Candidatus Heimdallarchaeota archaeon B3-JM-08 (PDB ID 3ORF) |
| **Data collection** |  |
| Beamline | SLS-X06DA |
| Wavelength (Å) | 0.999995 |
| Resolution range (Å) | 32.55 - 2.40 (2.489 - 2.403) |
| Space group | I 422 |
| Unit-cell parameters<br>a, b, c (Å)<br>$\alpha$, $\beta$, $\gamma$ (°) | a=34.193 b=34.193 c= 260.427<br>$\alpha=\beta=\gamma=90.00$ |
| Unique reflections | 5795 (930) |
| Multiplicity | 11.50 (11.74) |
| Completeness (%) | 99.74 (99.7) |
| Mean $I/\sigma(I)$ | 21.81 (6.90) |
| Wilson B-factor (Å<sup>2</sup>) | 31.69 |
| Rmerge | 0.071 (0.325) |
| CC1/2 | 0.999 (0.978) |
| Anisotropy | 0.177 |
| **Refinement** |  |
| Reflections used for refinement | 5782 |
| Reflections used for Rfree | 292 |
| Rwork | 0.2611 (0.3294) |
| Rfree | 0.2738 (0.3775) |
| **Model** |  |
| Protein atoms | 466 |
| Water molecules | 13 |
| Protein residues | 59 |
| **Average B-factors (Å<sup>2</sup>)** |  |
| Protein | 49.63 |
| Water | 47.3 |
| **Geometry** |  |
| R.m.s.d. bond lengths (Å) | 0.015 |
| R.m.s.d. bond angles (°) | 1.34 |
| **Validation** |  |
| Ramachandran favored (%) | 100 |
| Ramachandran outliers (%) | 0 |
| Rotamer outliers (%) | 3.64 |
| Clashscore | 3.12 |

The HPS1 structure reveals a parallel, left-handed α-helical coiled coil composed of four extended helices that wrap around one another in a highly symmetric arrangement (Fig. 2A). Each chain adopts an extended α-helical conformation, forming a tightly packed four-helix bundle. The core comprises 16 heptad layers of predominantly hydrophobic residues, numbered from −7 at the N-terminus to +8 at the C-terminus, with the central layer designated layer 0 (Fig. 2B).

**Figure 2.**
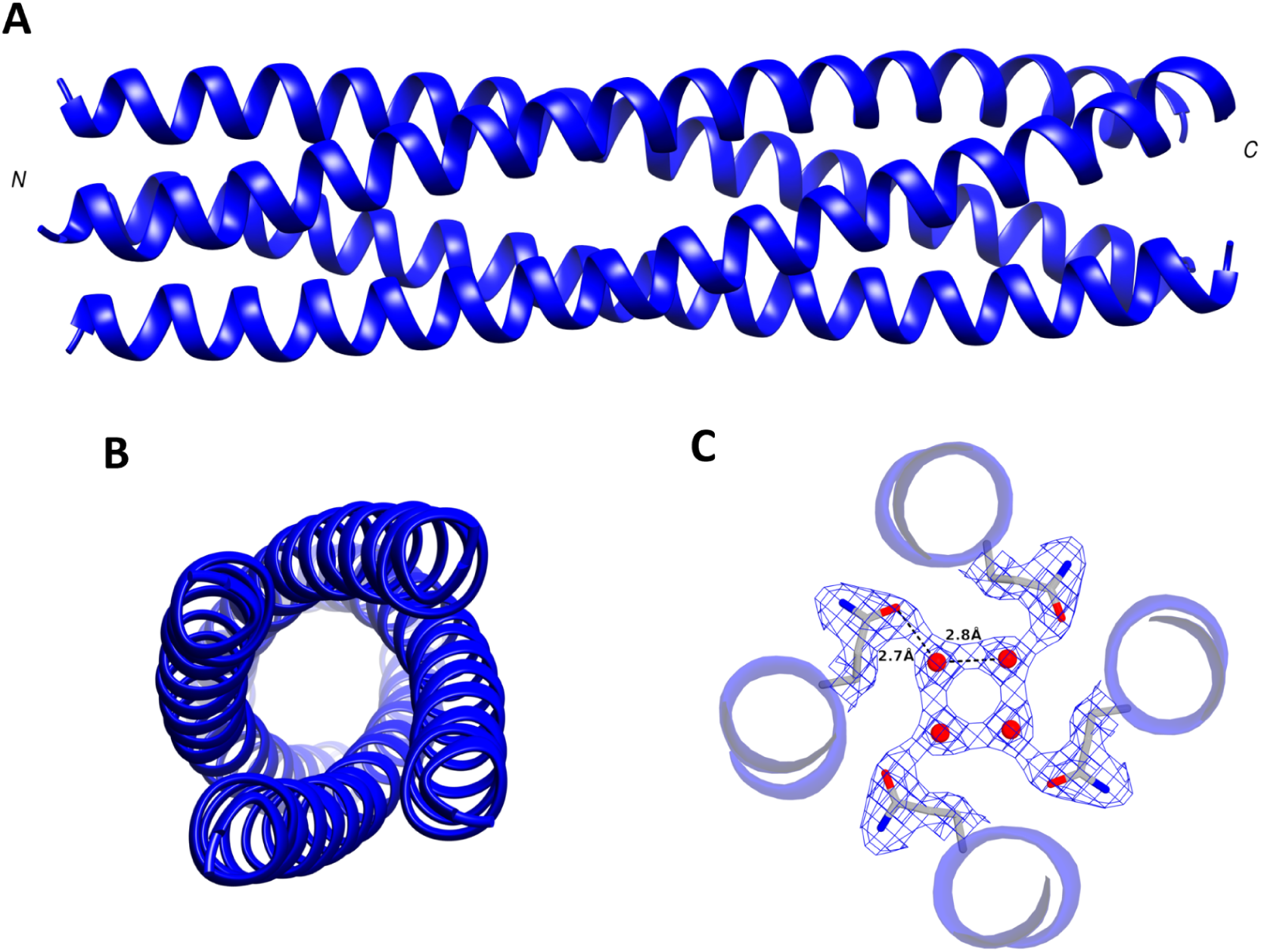
Crystal structure of the HPS1 homotetrameric SNARE-like complex. **(A)** Side view of the elongated HPS1 four-helix bundle, showing the symmetric arrangement of the four helices. **(B)** N-terminal view of the complex, illustrating its fourfold symmetry. **(C)** Close-up of the 0-layer, showing the four Q183 residues and the electron density surrounding the central region of the bundle. The Q183 side chains form a central polar layer, with water molecules occupying the cavity between them. Distances between the interacting residues and water molecules are indicated. The structures of selected hydrophobic layers are shown in Fig. S3.

A striking feature is the central polar layer (layer 0), formed by four conserved glutamine residues, Q183 (Fig. 2C). The four side chains point toward the center of the bundle and adopt similar conformations, generating a symmetric hydrogen-bonding network. A water-filled cavity at the center of the bundle further stabilizes this polar layer, with water molecules positioned to mediate hydrogen-bonding interactions between the glutamine side chains and backbone atoms at distances of approximately 2.7 Å.

Together, the structure reveals a highly symmetric homotetrameric coiled coil with a central glutamine layer and a hydrophobic core characteristic of SNARE-like four-helix bundles.

### Heterologous expression of HPS1 reveals ER localization in mammalian cells

Investigating the native localization and function of SNARE-like proteins in *Heimdallarchaeota* remains challenging because, although representatives of this lineage have recently been cultivated, no cultured strains were available for the experiments in this study. We therefore asked whether the Heimdall SNARE HPS1 can be heterologously expressed in mammalian cells and whether it associates with intracellular membranes. We cloned full-length HPS1, including its predicted C-terminal transmembrane domain, into a mammalian expression vector with an N-terminal enhanced green fluorescent protein (eGFP) fusion. The construct was transiently expressed in HeLa cells and human fibroblasts, and HPS1 localization was examined by fluorescence microscopy (Fig. 3). In both cell types, eGFP fluorescence was readily detectable and displayed a distinct reticular pattern throughout the cytoplasm, consistent with membrane association. To determine its subcellular localization, we co-expressed eGFP-HPS1 with the ER marker mCherry-SEC61. The green and red fluorescence signals showed substantial spatial overlap in both HeLa cells and fibroblasts, resulting in a yellow reticular pattern in merged images (Fig. 3). These observations are consistent with localization of HPS1 to the endoplasmic reticulum. Thus, the full-length Heimdall SNARE, including its C-terminal transmembrane domain, can be expressed in mammalian cells and is targeted to the ER. This finding demonstrates that HPS1 is compatible with the cellular machinery required for membrane insertion and ER localization in a eukaryotic cell.

**Figure 3.**
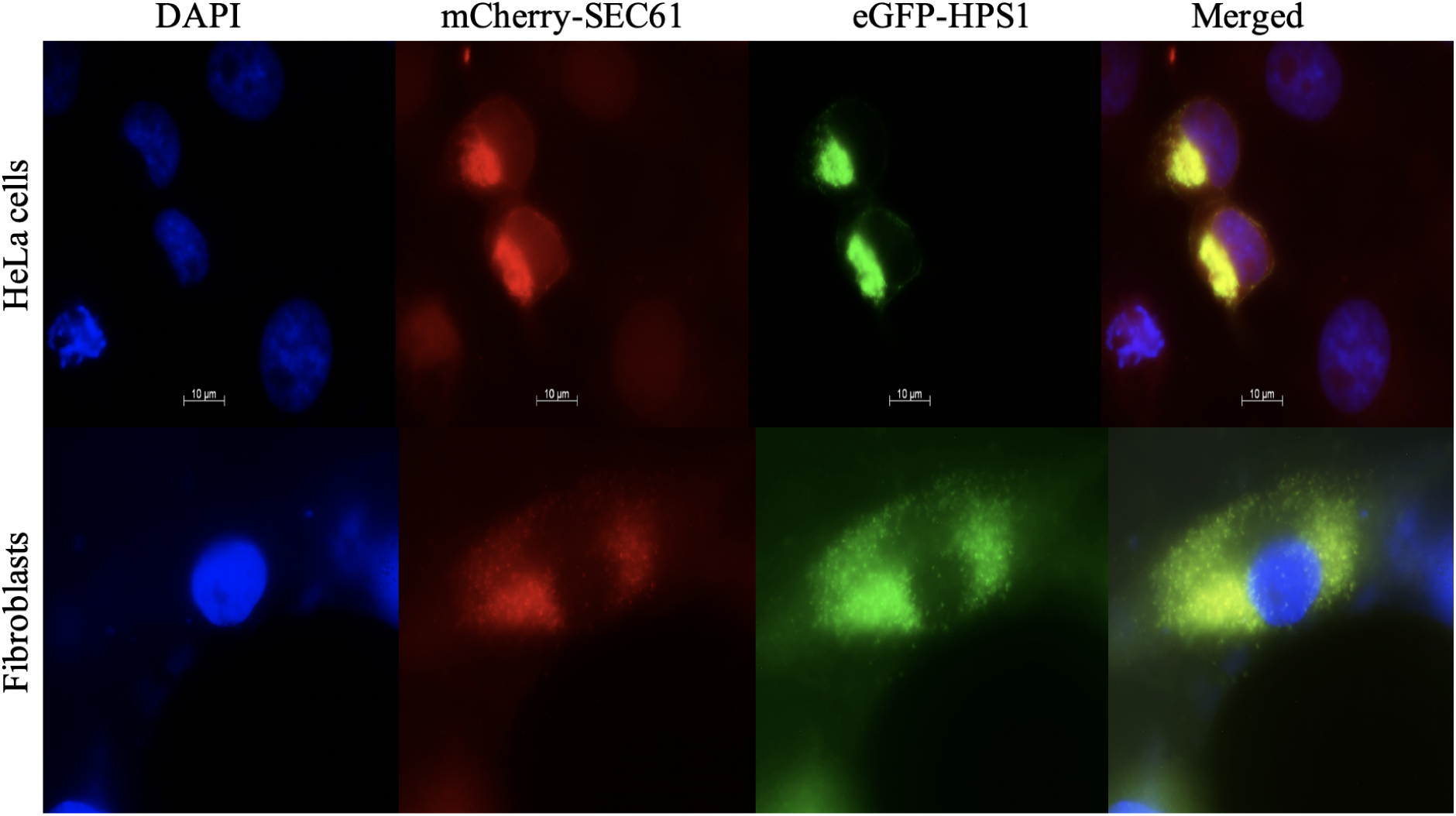
HPS1 localizes to the endoplasmic reticulum in mammalian cells. Fluorescence microscopy images of HeLa cells (top) and human fibroblasts (bottom) expressing eGFP-HPS1 (green) and the ER marker mCherry-SEC61 (red). The overlapping fluorescence signals appear yellow in the merged images, consistent with localization of HPS1 to the endoplasmic reticulum. Nuclei were stained with DAPI (blue). Scale bar, 10 µm.

### Heimdall SNAREs form a distinct clade separate from eukaryotic SNARE subtypes

In our earlier work, we identified four SNARE-like proteins in *Heimdallarchaeota*. With the continued growth of metagenomic sequence databases, we have now expanded this set to about 50 SNARE candidates. All were found in Heimdallarchaeia, including the orders *Heimdallarchaeaceae*, *Kariarchaeaceae*, and *Hodarchaeales*, but not in other Asgard lineages such as *Lokiarchaeia*, *Thorarchaeia*, or *Odinarchaeia*. Although most of the metagenome-assembled genomes (MAGs) are likely incomplete, it is notable that we detect with two exceptions (GCA_027018025 and GCA_027042105), in which a duplication occurred, usually a single SNARE-like gene per species in the available data.

All Heimdall SNARE candidates share a conserved domain architecture: a central SNARE motif flanked by a predicted C-terminal transmembrane domain and an N-terminal extension likely forming a three-helix bundle. This arrangement resembles the general domain structure found in many eukaryotic SNAREs. To assess their evolutionary relationships, we constructed a phylogenetic tree based on a multiple sequence alignment of SNARE motifs from both Heimdall and eukaryotic SNAREs from 24 different species as we did in our previous study ^31^. The resulting tree (Fig. 4A) shows that all Heimdall SNARE sequences group into a well-supported monophyletic clade, clearly separate from the four canonical eukaryotic SNARE types: Qa-, Qb-, Qc-, and R-SNAREs, which correspond to the four defined positions within the eukaryotic four-helix SNARE complex. In addition, we constructed a phylogenetic tree of all Heimdall SNAREs identified to date (Fig. 4B) and visualized the conservation pattern of their SNARE motifs as a web logo (Fig. 4C).

**Figure 4:**
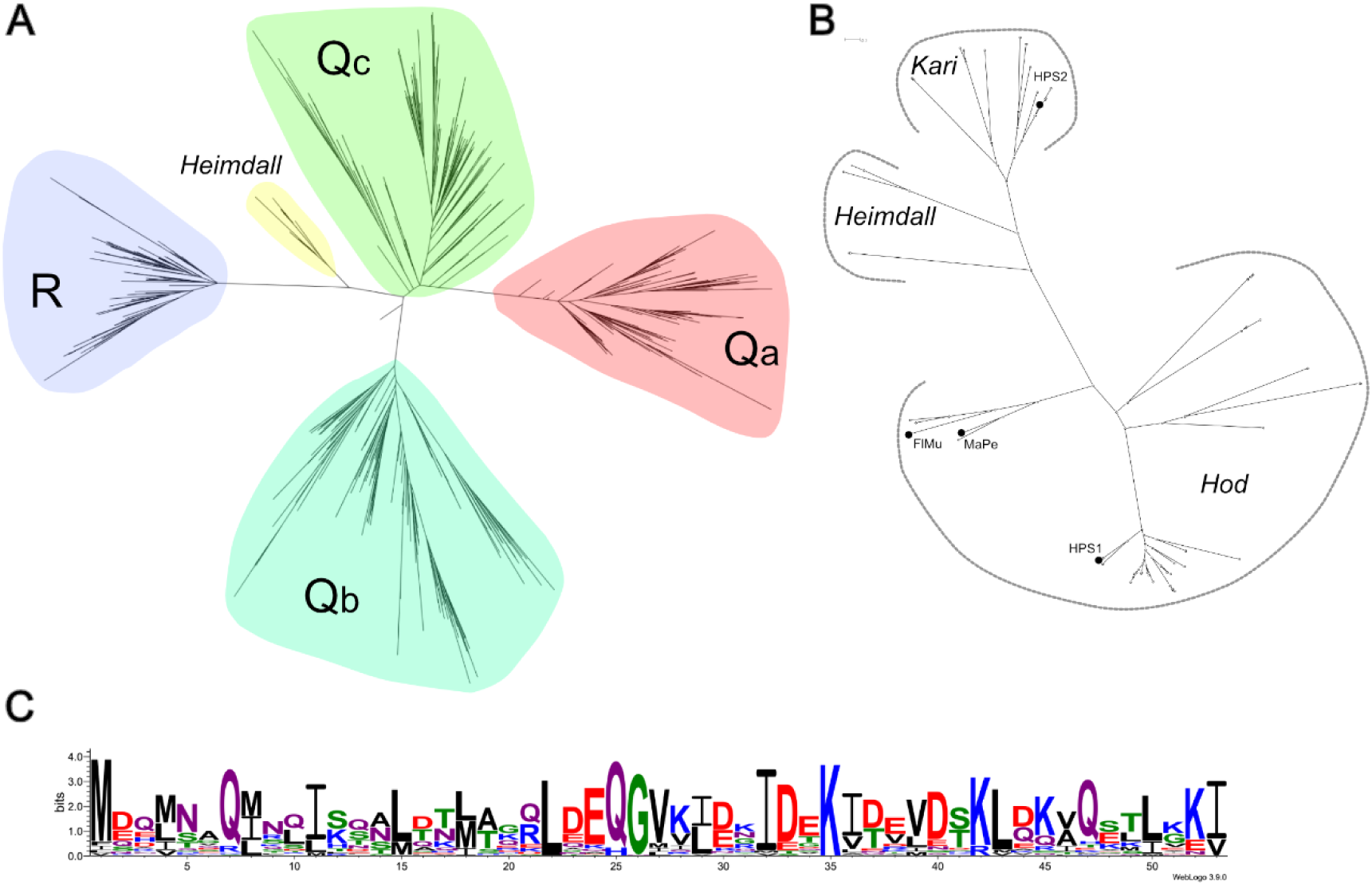
Phylogenetic relationships of Heimdall SNARE proteins. **(A)** Maximum-likelihood phylogenetic tree of the SNARE motifs from 24 representative eukaryotic species and 20 Heimdall SNARE proteins. The tree is based on an alignment of the SNARE motif sequences only and is adapted from Neveu et al. 2020 ^31^. Eukaryotic SNAREs segregate into the four conserved subfamilies, **Qa**, **Qb**, **Qc**, and **R**, whereas the Heimdall SNAREs form a distinct branch basal to the eukaryotic SNAREs. **(B)** Maximum-likelihood phylogenetic tree of approximately 52 Heimdall SNARE proteins inferred from an alignment of their full-length amino acid sequences. Three major clades, labeled Kari, Hod, and Heimdall, are highlighted for clarity. These designations are used to distinguish the principal groups represented in the tree and do not imply a specific taxonomic rank, as the classification of Asgard archaea remains under active revision. The two proteins characterized in this study, HPS1 and HPS2, are indicated; HPS1 was used for X-ray crystallographic structure determination. SNARE proteins from the recently cultivated Asgard archaea *Ca.* ‘Flexarchaeum multiprotrusionis’ (FlMu) and ‘Margulisarchaeum peptidophila’ (MaPa) are also indicated. **(C)** Sequence conservation of the Heimdall SNARE motif displayed as a WebLogo representation based on the alignment of all Heimdall SNARE sequences. Residue height is proportional to the degree of conservation at each position. All identified Heimdall SNARE sequences are available in the Tracey database (https://tracey.unil.ch)^40^, which provides sequence identifiers and annotations of the SNARE motif and N-terminal region. A Heimdall SNARE-specific hidden Markov model (HMM) was generated to identify and delineate the SNARE motif in these sequences.

The distinct phylogenetic placement of Heimdall SNAREs suggests that they represent a lineage that diverged before the expansion and specialization of SNAREs in eukaryotes. The presence of only a single SNARE gene per genome further supports the idea that the last common ancestor of eukaryotic SNAREs may have been a homomeric system with limited functional diversity. These findings are consistent with a scenario in which eukaryotic SNARE complexity arose from an ancestral, possibly Heimdall-like, SNARE toolkit through gene duplication and functional divergence.

### Molecular dynamics reveal conserved and divergent features of Heimdall SNARE architecture

To investigate whether the Q/R asymmetry characteristic of eukaryotic SNARE complexes ^32,41^ is sufficient to establish the canonical 0-layer, we performed molecular dynamics simulations of the wild-type Heimdall SNARE complex and a mutant in which one of the four 0-layer glutamines was replaced by arginine. In the wild-type complex, the four glutamines formed a compact polar layer (Fig. 2C), with Gln–Gln hydrogen-bond contacts occurring frequently throughout the simulations (0.30 contacts per frame on average). In contrast, the engineered Arg183 did not insert into the center of the bundle or form the characteristic buried interactions with the three remaining glutamines; Arg–Gln hydrogen-bonding was essentially absent (0.016 contacts per frame), and the guanidinium group consistently pointed outward from the bundle axis in all five mutant simulations. The substitution caused a modest local splaying of the N-terminal ends of the bundle (12.5 versus 10.0 Å) but did not compromise its overall stability. Thus, introducing an arginine into the Heimdall 0-layer does not recreate the buried ionic layer characteristic of eukaryotic SNARE complexes. Both the wild-type and R3Q bundles remained stable and folded across all five 100 ns replicas, with low core backbone RMSD (1.38 ± 0.09 and 1.28 ± 0.04 Å), indistinguishable core radii of hydration (24.3 Å for both), and comparable per-residue Cα fluctuations (RMSF 0.88 ± 0.02 vs 0.85 ± 0.01 Å), confirming that the arginine substitution does not destabilize the bundle (Fig. S5). Rather, the native 4Q architecture appears structurally adapted to a compact, uncharged polar layer, with the bulky charged arginine accommodated by displacement toward the solvent-exposed surface. These results suggest that the eukaryotic Q/R 0-layer was not established simply by replacing one glutamine with arginine, but emerged together with the broader asymmetric architecture of the eukaryotic SNARE bundle.

To further compare the architecture of the Heimdall and eukaryotic SNARE bundles, we quantified the volume of their hydrophobic cores using the cubicle method described by Yadav et al. ^32^. This approach calculates the convex-hull volume defined by the Cα atoms of the eight core positions contributed by four adjacent a/d layers and follows the volume along the length of the bundle. Applying the same automated register-walk procedure to the Heimdall MD ensembles and five representative eukaryotic SNARE crystal structures revealed that the Heimdall bundle has a core volume within the range observed for eukaryotic SNARE complexes (Fig. S6). Moreover, the Heimdall bundle reproduced the characteristic expansion of the core toward the center of the bundle, resulting in a similar spindle-shaped profile. Thus, despite differences in sequence and in the molecular organization of the 0-layer, the overall core packing of the Heimdall four-helix bundle closely resembles that of eukaryotic SNARE complexes.

## Discussion

Our structural analysis reveals that SNARE-like proteins from Heimdallarchaea assemble into highly stable homotetrameric four-helix bundles that are strikingly similar to canonical eukaryotic SNARE complexes. This structural resemblance, despite the homomeric nature of the Heimdallarchaeal complex, suggests that the fundamental mechanism of SNARE-mediated membrane fusion may have originated before the emergence of the last eukaryotic common ancestor (LECA). In contrast to eukaryotic SNARE complexes—composed of four distinct helices contributed by Qa-, Qb-, Qc-, and R-SNAREs—the Heimdallarchaeal complex is formed by four identical helices. This homotetrameric assembly could represent a functional or pre-functional ancestor of the eukaryotic SNARE fusion machinery.

Although the fusion activity of Heimdallarchaeal SNAREs has not yet been demonstrated in vivo, the structural integrity and thermostability of the complex are consistent with a membrane-fusogenic role. The presence of intracellular vesicle-like structures in recently cultivated Heimdallarchaeota ^23,24^ is compatible with this interpretation. In contrast, such compartments have not been observed in cultivated Lokiarchaeota ^21,22^, suggesting that vesicle-based compartmentalization may have emerged or become elaborated specifically within the Heimdall lineage. Consistent with this scenario, SNARE-like genes are thus far restricted to Heimdallarchaeota and are conserved across its currently recognized subgroups ^42,43^, in line with proposals that Heimdallarchaeota represent the closest archaeal sister group to eukaryotes.

Additional components of membrane trafficking are also present in Asgard archaea. In particular, the identification of Sec23- and Sar1-like proteins indicates the presence of components functionally analogous to the COPII system. Moreover, additional endomembrane-associated features, including elements of the oligosaccharyltransferase complex ^12^; ^44^, are already present in Asgard genomes. Together with the presence of SNAREs and vesicle-like structures in Heimdallarchaeota, these observations are consistent with the existence of a rudimentary membrane-remodeling and trafficking system in the archaeal lineage leading to eukaryotes. Such a system could have provided a cellular foundation upon which the more elaborate eukaryotic endomembrane system subsequently evolved.

The evolutionary implications of this scenario extend beyond individual membrane-trafficking proteins. Eukaryotic genomes are chimeric, reflecting the evolutionary merger of an archaeal host and a bacterial endosymbiont ^45^. The archaeal contribution includes the core genetic information system, encompassing the machineries for DNA replication ^46^, transcription, and translation, which are phylogenetically and mechanistically archaeal in nature ^47^, but also a substantial fraction of the conserved cellular machinery characteristic of eukaryotes. In contrast, genes of bacterial origin are largely associated with bioenergetic and mitochondrial functions and derive predominantly from the alphaproteobacterial ancestor of mitochondria, whereas additional bacterial contributions appear to reflect more sporadic horizontal gene transfers rather than a second major founding lineage ^48^; ^7,49^; ^44,50–52^.

The growing body of evidence from Asgard archaea indicates that the archaeal host was not a simple prokaryotic cell but already possessed a diverse repertoire of cellular machinery. Asgard genomes encode numerous eukaryotic signature proteins (ESPs), including actin, tubulins, small GTPases, and components of the ESCRT machinery ^11–14^. Yet these systems generally appear considerably less diversified than their eukaryotic counterparts. In the ESCRT-III system, for example, an ancestral duplication in Asgard archaea has been proposed to have generated the precursors of the CHMP1-3 and CHMP4-7 subfamilies in eukaryotes ^25–27^. Such examples suggest that eukaryotic cellular complexity may have arisen not through the de novo invention of entirely new molecular systems, but through the duplication, diversification, and specialization of a simpler ancestral toolkit.

A similar pattern is evident in AAA+ ATPases associated with proteasome regulation and membrane trafficking. The archaeal proteasome-associated ATPase PAN forms a homohexameric ring, whereas the eukaryotic 19S regulatory particle contains six distinct AAA+ ATPase subunits, Rpt1–Rpt6, forming a heterohexameric ring ^53^. Archaeal Cdc48 contains two tandem AAA+ ATPase domains and has been proposed to have arisen through duplication of an ancestral ATPase such as PAN. In eukaryotes, Cdc48 subsequently diversified into several specialized paralogs, including NSF, which disassembles SNARE complexes following membrane fusion ^54^. We found that Heimdallarchaeal SNARE complexes are resistant to disassembly by murine NSF, raising the possibility that an archaeal Cdc48-like ATPase may have fulfilled an analogous function before the emergence of the specialized eukaryotic NSF machinery. This possibility remains to be tested experimentally.

The SNARE system itself illustrates this transition particularly clearly. The archaeal homomeric complex characterized here contrasts with the four distinct SNARE classes, Qa, Qb, Qc, and R, that form the canonical heterotetrameric eukaryotic complex. This transition from a homomeric to a heteromeric architecture may have enabled tighter and more specific packing of the four-helix bundle ^32^, followed by the expansion of specialized SNARE families associated with individual trafficking pathways in the LECA ^55^. Comparable elaborations occurred in other vesicle-associated systems, including Sec1/Munc18 (SM) proteins ^56^, CATCHR-family tethering factors ^57,58,59,60^, cytoskeletal motors ^61–67^, and membrane coat protein complexes such as COPI, COPII, and clathrin ^68^; ^69–71^. Many of these coat proteins are evolutionarily related to components of the nuclear pore complex and intraflagellar transport systems, suggesting that vesicle coats and other macromolecular trafficking assemblies share deeper evolutionary relationships ^72^; ^73^.

At the same time, canonical eukaryotic coat complexes, SM proteins, multisubunit tethering complexes, and cytoskeletal motor proteins have not been identified in Asgard genomes ^50,74^. This contrast is informative: prototypic elements of membrane remodeling and trafficking were already present in the archaeal lineage, whereas the extensive regulatory, mechanical, and spatial integration characteristic of the eukaryotic endomembrane system appears to have emerged later through gene duplication, diversification, and architectural innovation.

Our findings therefore support a model in which key elements of the eukaryotic endomembrane system, including the SNARE-mediated membrane-fusion machinery, have deep evolutionary roots in the archaeal lineage. The striking structural conservation of the SNARE four-helix bundle, despite its simpler homomeric organization and distinct 0-layer chemistry, provides a concrete example of how an archaeal molecular system could have served as a starting point for subsequent eukaryotic diversification. Together with the presence of membrane-associated compartments and additional trafficking components in Heimdallarchaeota, these findings support the view that the archaeal host of eukaryogenesis possessed a foundational membrane-remodeling toolkit that was subsequently expanded and specialized during the emergence of the eukaryotic cell.

## Methods

### Protein expression and purification

Codon-optimized genes encoding Heimdallarchaeal SNARE proteins (PWI47941.1 from Candidatus Heimdallarchaeota archaeon B3-JM-08, HPS1; and OLS22354.1 from Candidatus Heimdallarchaeota archaeon LC_2, HPS2) were synthesized and subcloned into the pET28a vector (GenScript) as described ^31^. To enable UV detection at 280 nm, a phenylalanine-to-tryptophan substitution was introduced at position 155 in HPS1 (HPS1^F155W). Proteins were expressed in E. coli BL21 (DE3) and purified by Ni²⁺-NTA affinity chromatography. Following thrombin-mediated removal of the His₆ tag, proteins were further purified by anion exchange chromatography (Mono Q HR 10/10, GE Healthcare) using an ÄKTA system. Elution was performed with a linear NaCl gradient in 20 mM Tris (pH 7.4), 1 mM EDTA, as described previously ^31^; ^75,76^. Purity (∼95%) was assessed by SDS–PAGE. Protein monodispersity was evaluated by size exclusion chromatography on a Superdex 200 HR 10/30 column. Protein concentrations were determined by absorbance at 280 nm and by Bradford assay.

### Size-exclusion chromatography coupled to multi-angle light scattering (SEC–MALS)

HPS1^F155W was prepared in 20 mM HEPES (pH 7.4), 150 mM NaCl and filtered (0.1 µm). A 100 µL sample (4 mg mL⁻¹) was applied to a Superdex 200 Increase 5/150 GL column equilibrated in the same buffer. Elution was performed at a flow rate of 0.25 ml min⁻¹. Light scattering was measured using a miniDAWN TREOS detector (Wyatt Technology), and molar masses were calculated using ASTRA 6 software.

### Circular dichroism (CD) spectroscopy

CD measurements were performed essentially as described ^31,75,76^ using a Chirascan spectrometer (Applied Photophysics) and quartz cuvettes (0.1 cm path length). Proteins were dialyzed against 20 mM sodium phosphate (pH 7.4), 150 mM NaCl. Far-UV CD spectra were recorded from 200–260 nm at a bandwidth of 1 nm. For thermal denaturation experiments, ellipticity at 220 nm was monitored between 10 and 100 °C at a heating rate of 30 °C h⁻¹. For chemical denaturation, proteins were incubated at room temperature for ∼5 h in the same buffer supplemented with increasing concentrations of urea or guanidinium hydrochloride (GdnHCl). Spectra were recorded from 210–260 nm to avoid interference from denaturant absorption below 210 nm.

### Crystallization and data collection

Initial crystallization screening was performed against ∼500 conditions at the Protein Production and Structure Core Facility (PPSCF, EPFL, https://www.epfl.ch/research/facilities/ptpsp/). Crystals were obtained by sitting-drop vapor diffusion by mixing equal volumes (0.2 µL) of protein solution (13 mg mL⁻¹) and reservoir solution. Drops were equilibrated against 20 µL reservoir solution in 96-well plates using a Mosquito liquid handling robot (SPT Labtech) and incubated at 20 °C. Crystals appeared after several days in a condition containing 0.1 M ammonium acetate, 0.1 M zinc chloride, 15% (v/v) PEG Smear High, and 0.1 M Bis-Tris–HCl (pH 7.2). Crystals were cryoprotected with reservoir solution supplemented with 25% glycerol and flash-cooled in liquid nitrogen. Diffraction data were collected at the Swiss Light Source.

### Structure determination and refinement

The structure of the HPS1 SNARE-like homotetramer was solved by molecular replacement using PHENIX (v.1.20.1) (Adams et al., 2011), with Syntaxin-6 (PDB ID: 2NPS, chain D) as search model. The initial model was generated using MoRDa (Krissinel et al., 2018). Iterative model building and refinement were carried out using PHENIX (phenix.refine) and Coot (v.0.9.8) (Casañal et al., 2020). Structural figures were prepared using PyMol.

### Cell culture and transfection

An N-terminal eGFP-tagged HPS1 construct was generated in the pcDNA3.1(+)-N-eGFP vector (GenScript). Plasmid DNA was amplified in E. coli NEB 5-alpha and purified using a QIAGEN Plasmid Midi Kit. DNA concentration and purity were assessed spectrophotometrically.

HeLa cells and fibroblasts were seeded at 2 × 10⁵ cells per well in 12-well plates and cultured in DMEM supplemented with 10% fetal bovine serum. Transfection was performed after 24 h using FuGENE (Promega) at a 3:1 reagent-to-DNA ratio. Cells were fixed 20 h post-transfection in 4% paraformaldehyde in PBS for 10 min at room temperature and washed with PBS. For nuclear staining, cells were incubated with DAPI (1:10,000) for 10 min, washed, and mounted in Mowiol 4-88. Alternatively, samples were mounted in Vectashield. Slides were incubated overnight at room temperature prior to fluorescence imaging using an AxioVision system.

### Phylogenetic analysis of Heimdall SNARE proteins

#### Sequence collection and identification of SNARE proteins

Asgard archaeal genome assemblies and their predicted protein sequences were retrieved from the NCBI database on September 22, 2025. Approximately 1,000 Asgard archaeal metagenome-assembled genomes (MAGs) were available at the time of data collection. The corresponding predicted protein sequences were downloaded and screened for SNARE proteins using hidden Markov models (HMMs) developed by Kloepper et al. ^55^ and the hmmsearch program of HMMER ^77^; ^78^. An E-value threshold of 10^−4^ was used for the HMM searches. Candidate SNARE proteins were retained for subsequent phylogenetic analyses when a SNARE motif could be reliably identified.

For the identification and annotation of Heimdall SNAREs, a Heimdall SNARE-specific HMM was generated from the identified sequences and used to define the boundaries of the SNARE motif. The resulting sequences, together with their sequence identifiers and annotations of the SNARE motif and N-terminal region, were deposited in the Tracey database (https://tracey.unil.ch) ^40^.

#### Phylogenetic analysis of full-length Heimdall SNAREs

For analysis of the evolutionary relationships among Heimdall SNAREs, full-length protein sequences were aligned using MUSCLE ^79^ with default parameters. Sequences for which the alignment of the SNARE region was considered unreliable were excluded. Maximum-likelihood phylogenetic trees were inferred using IQ-TREE2 ^80^ under the LG+I+G4 substitution model, and branch support was assessed using 1,000 bootstrap replicates.

#### Phylogenetic relationship between Heimdall and eukaryotic SNAREs

To place Heimdall SNAREs in the context of eukaryotic SNARE evolution, we constructed a separate maximum-likelihood phylogenetic tree using SNARE proteins from 24 representative eukaryotic species previously analyzed by Neveu et al. ^31^, together with 20 Heimdall SNARE proteins. Only the SNARE motif was considered for this analysis. The 53-residue SNARE motifs were identified using HMMER v3.2.1 and used directly for phylogenetic reconstruction following the procedure described ^31^. Because the motif boundaries were defined by the HMM, no additional global sequence alignment was required ^55^. Phylogenetic reconstruction was performed using IQ-TREE2 as described above, with branch support assessed from 1,000 bootstrap replicates.

### Molecular Dynamics Simulations

The asgard SNARE four-helix bundle was studied as a homotetramer (chains A-D; residues 157-215). Two systems were prepared: the WT bundle, in which the 0-layer, residue 183, is formed by four glutamines (4Q), and an R3Q 0-layer variant in which Gln183 of chain A was mutated to an arginine to mimic the eukaryotic R-Q-Q-Q 0-layer geometry.

Both systems were built with CHARMM-GUI ^81^ using the CHARMM36m all-atom protein force field ^82^. Each bundle was solvated in an orthorhombic box of TIP3P water ^83^ with a minimum solvent padding sufficient to give box dimensions of approximately 110 x 104 x 90 Å. The systems were neutralized with Na and Cl ions and brought to a physiological ionic strength of 0.15 M.

All simulations were performed with NAMD3 ^84^. The time step for integrations was 2 fs. Short-range non-bonded interactions were computed with a 12 Å cutoff and a force-based switching function applied from 10 Å, with a 16 Å pair-list distance. Long-range electrostatics were treated with the particle-mesh Ewald (PME) method ^85^. Periodic boundary conditions were applied in all three dimensions.

Each system was first minimized for 10000 steps and then equilibrated for 250 ps with harmonic positional restraints on the protein heavy atoms, during which velocities were reassigned to the target temperature every 500 steps. Production simulations were run in an isothermal-isobaric (NPT) ensemble at 303.15 K and 1.01325 bar. Temperature was maintained by a Langevin thermostat with a 1 ps⁻¹ damping coefficient, and pressure was controlled with a Nosé-Hoover Langevin piston barostat ^86^; ^87^ with a 50 fs oscillation period and 25 fs decay time.

For each system, five independent replicas were run for 100 ns each, differing in their randomized initial velocities, giving 500 ns of sampling per system. Coordinates were saved every 100 ps, yielding 1000 frames per replica.

#### Trajectory analysis

All analyses were performed with MDAnalysis ^88^; ^89^ over the full 100 ns of each replica; equilibration was confirmed by verifying that the first and last 50 ns means of the core backbone RMSD and radius of gyration agreed to within ∼0.1 Å per replica. Structural metrics were computed on the Cα atoms of residues 160-213, after aligning all frames to the first frame. Reported quantities include the backbone RMSD, the core radius of gyration, per-residue Cα RMSF, secondary structure content (DSSP), and inter-chain center-of-mass distances. Per-replica values are reported as means over the trajectory, and cross-replica statistics as mean ± standard deviation over the five replicas (n = 5).

#### Inner volume analysis

The inner volume of the four-helix bundle was quantified following the virtual cubicle approach of Yadav et al. ^32^. For each core layer, the available volume was computed from the cubicle defined by the Cα atoms of the four chains at the ‘a’ and ‘d’ heptad positions, and a per-layer profile was built along the bundle. The identical estimator was applied to the Asgard MD ensembles (averaged over all frames of the five replicas) and to five eukaryotic SNARE crystal structures (PDB ids: 1SFC, 2NPS, 3B5N, 1GL2 and 4WY4), each registered on its own R-Q-Q-Q 0-layer, to compare the Asgard core packing to the conserved eukaryotic spindle.

## Acknowledgment

This work was supported by the Swiss National Science Foundation (Grant 310030_219549 to D.F.). We thank Jacqueline Chrast for her assistance with mammalian cell culture and staining experiments. We are grateful to Florence Pojer and Kelvin Lau at the EPFL Protein Production and Structure Platform (PTPSP) for their support with protein crystallization and X-ray structure determination. We also thank all members of the Fasshauer laboratory for their helpful discussions, technical assistance, and for creating a stimulating and enjoyable working environment.

## Author contributions

Iman Bentahar: Conceptualization; investigation; methodology; validation; writing – review and editing; data curation. Deepak Yadav: Conceptualization; investigation; methodology; validation; writing – review and editing; data curation. Aysima Hacisuleyman: Conceptualization; investigation; methodology; validation; writing – review and editing; data curation. Dirk Fasshauer: Conceptualization; funding acquisition; writing – original draft; writing – review and editing; methodology; validation; visualization; supervision; resources.

## Data availability statement

The data that support the findings of this study are openly available in the Zenodo repository 10.5281/zenodo.23390284.

## Abbreviations

(SNARE): Soluble N-ethylmaleimide-sensitive factor Attachment protein REceptor
(NSF): N-ethylmaleimide-sensitive factor
(ER): endoplasmic reticulum
(LECA): Last Eukaryotic Common Ancestor
(MD): molecular dynamics
(HMMs): hidden Markov models
(PAGE): polyacrylamide gel electrophoresis
(RMSD): Root Mean Square Deviation
(ESPs): eukaryotic signature proteins
(SEC): size-exclusion chromatography
(CD): circular dichroism spectroscopy
(GdnHCl): guanidinium hydrochloride
(HPS): Heimdallarchaeota Prototypic SNARE
(PCA): principal component analysis

## Supplements

**Figure S1.**
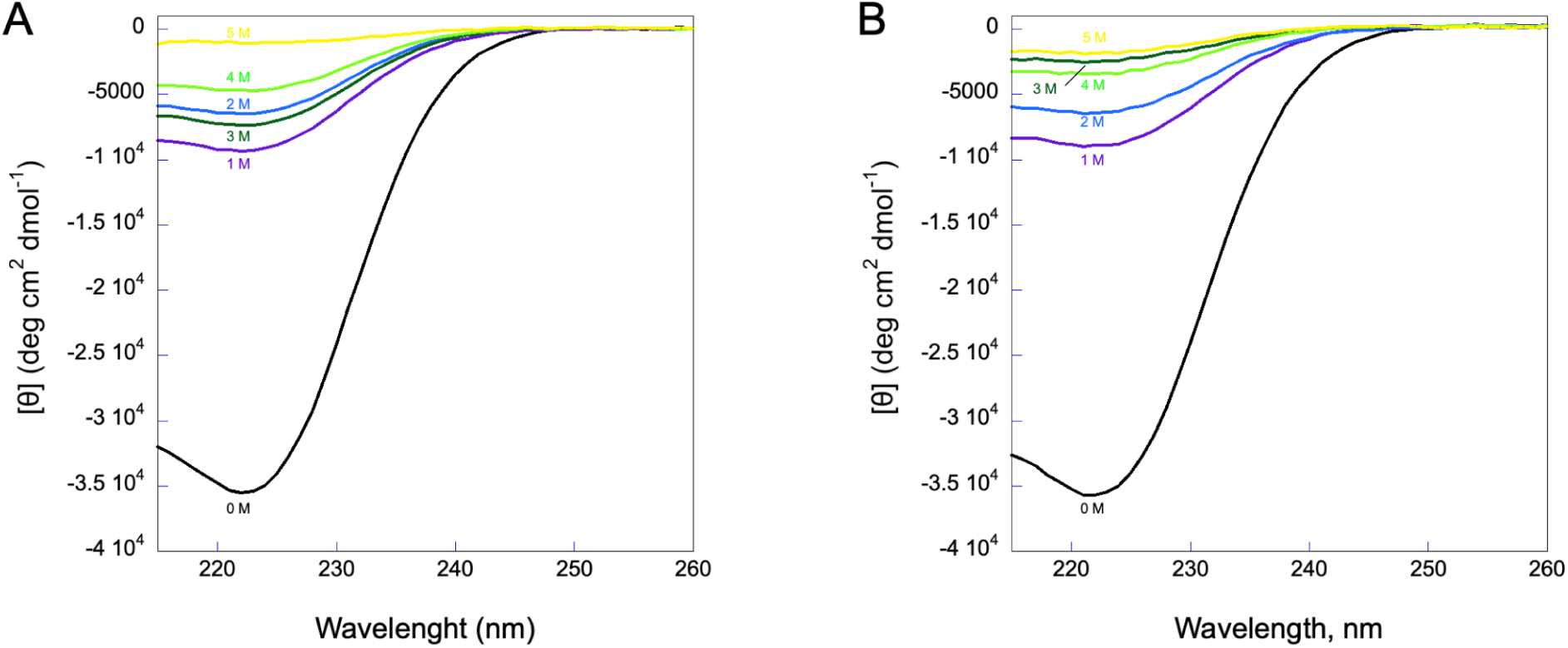
Chemical denaturation of Heimdall SNARE-like proteins monitored by CD spectroscopy. (A) CD spectra of HPS1 recorded in the presence of increasing concentrations of GdnHCl (1–5 M). (B) CD spectra of HPS2 recorded under the same conditions. Spectra in the absence of GdnHCl are shown as controls. Proteins were prepared at final concentrations of 23.6 µM (HPS1) and 23.9 µM (HPS2) in 20 mM Na₃PO₄ (pH 7.4), 150 mM NaCl and pre-incubated with GdnHCl for 5 h at room temperature. CD spectra were recorded from 210–260 nm to minimize interference from GdnHCl absorption below 210 nm.

**Figure S2:**
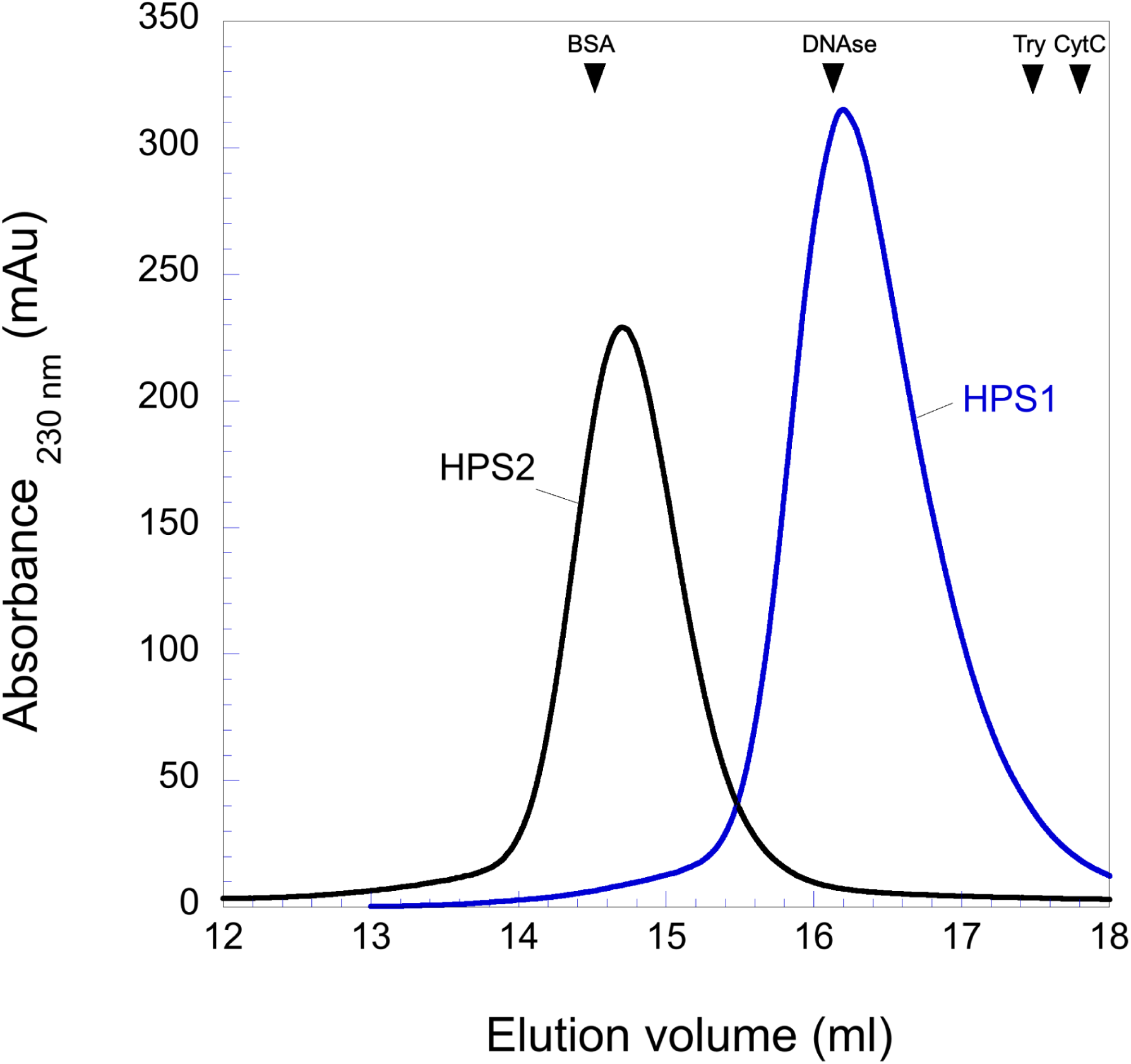
HPS1 and HPS2 proteins form homomeric complexes. The elution profiles of HPS1 and HPS2 proteins were monitored at 230 nm. The experiments were conducted using a Superdex^TM^ 200 10/300 GL column equilibrated in a buffer containing 20 mM Tris-HCl (pH 7.4) and 150 mM NaCl with a flow rate of 0.5 mL/min. The elution volumes of calibration proteins are indicated above.

**Figure S3:**
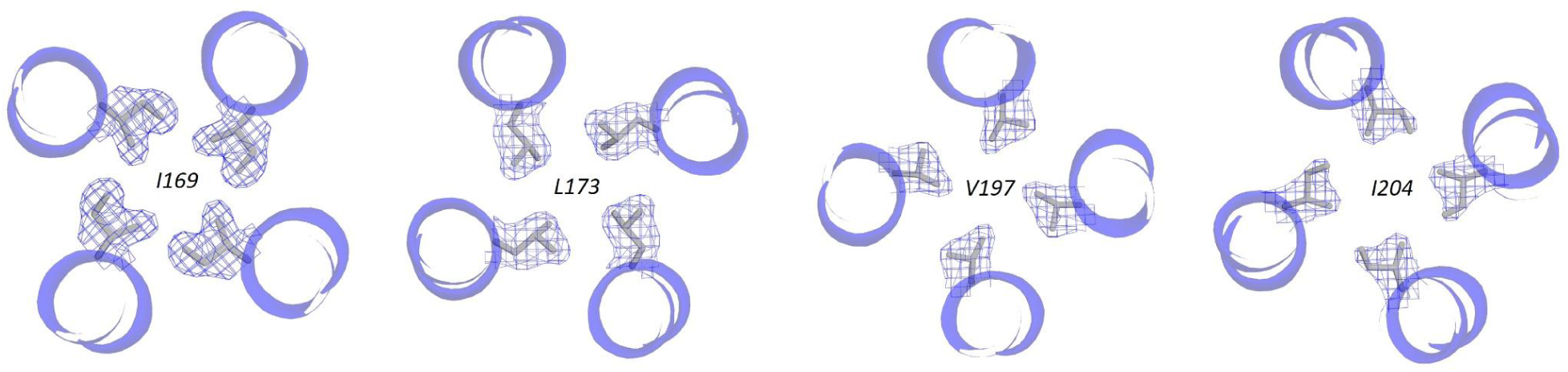
Intermolecular contacts within the hydrophobic core of the HPS1 complex. N-terminal views of four representative hydrophobic layers of the HPS1 four-helix bundle, corresponding to Ile169 (−4), Leu173 (−3), Val197 (+4), and Ile204 (+6). The views illustrate the fourfold-symmetric arrangement and packing of the side chains around the bundle axis. The four α-helices are shown as blue cartoons, with the selected hydrophobic side chains displayed as sticks (carbon atoms in grey). The corresponding electron density is shown as a blue mesh.

**Figure S4:**
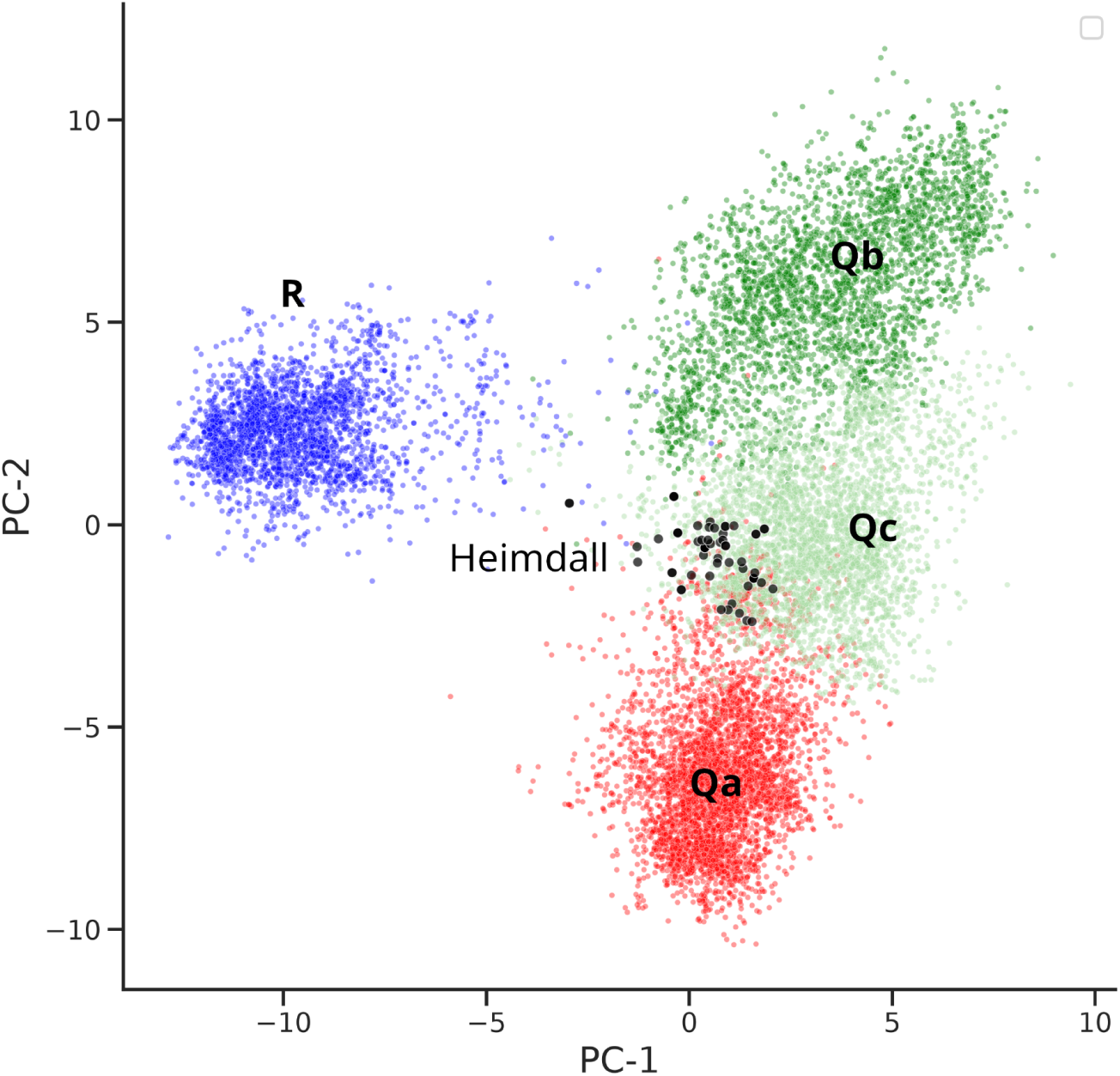
Heimdall SNAREs can be distinguished from the four eukaryotic SNARE types based on amino acid composition. Principal component analysis (PCA) of SNARE amino acid compositions was performed using six physicochemical descriptors of amino acid side chains: hydrophobicity, van der Waals volume, flexibility, bulkiness, steric factor, and maximum number of hydrogen bonds. The first two principal components are shown. In our previous study, we showed that the four eukaryotic SNARE types can be distinguished in this way ^32^. Heimdall SNAREs occupy a distinct region of the PCA space and can be distinguished from the four eukaryotic SNARE types based on these physicochemical properties. R-SNAREs are shown in blue, Qa-SNAREs in red, and Qb- and Qc-SNAREs in dark and light green, respectively. Heimdall SNAREs show the greatest overlap with Qc-SNAREs, which also represent the least divergent eukaryotic SNARE group in the phylogenetic analysis.

**Figure S5.**
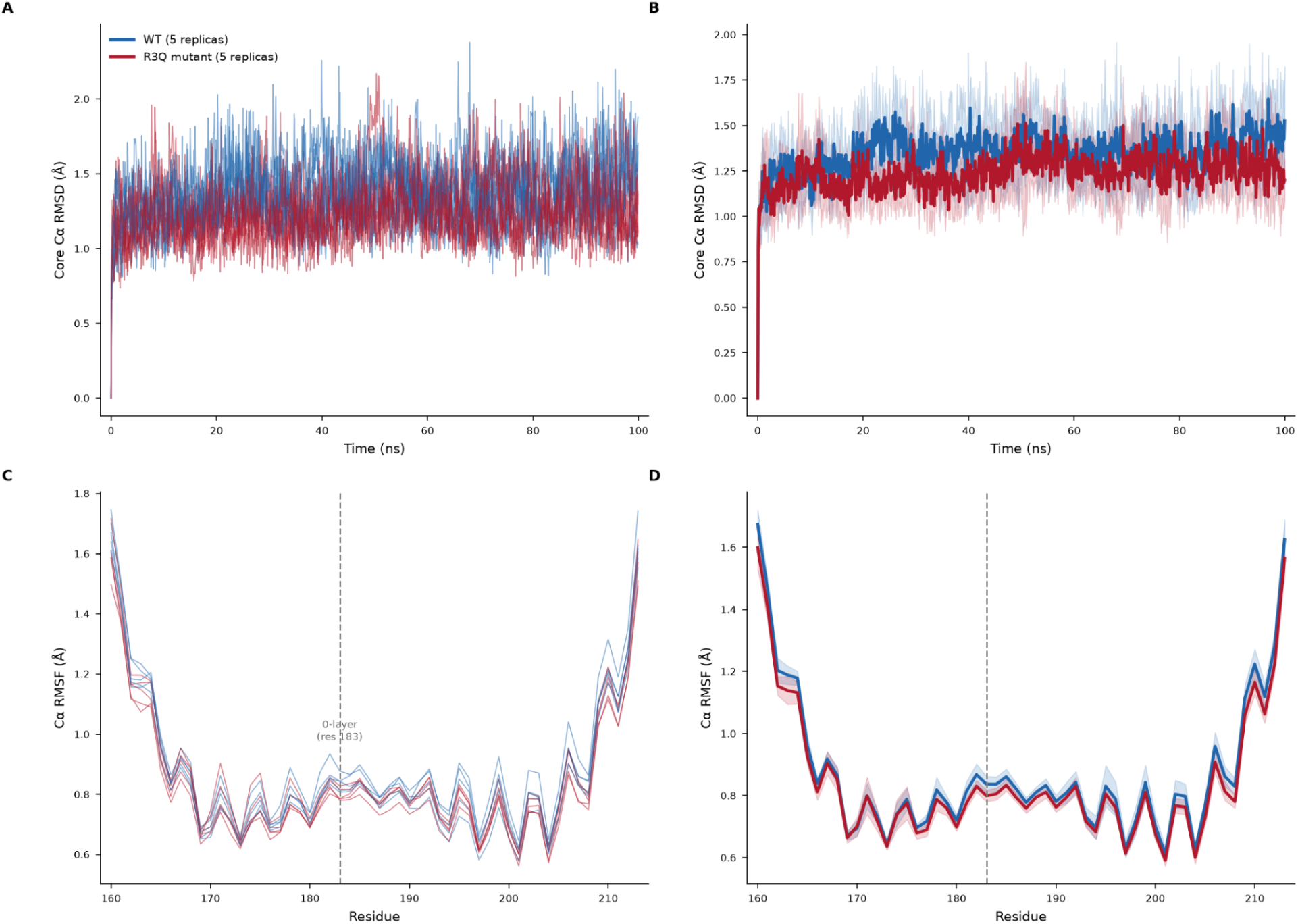
R3Q does not destabilize the bundle. Core Cα RMSD (panels a,b) and per-residue RMSF (panels c,d) for wild-type (blue) and R3Q 0-layer mutant (red) over 100 ns replicas each; colors in all panels follow the legend in panel a. (a,c) Individual replicas overlaid; (b,d) across-replica mean (bold lines) ± 1 SD (shaded). Dashed line in panels c and d marks the 0-layer.

**Figure S6:**
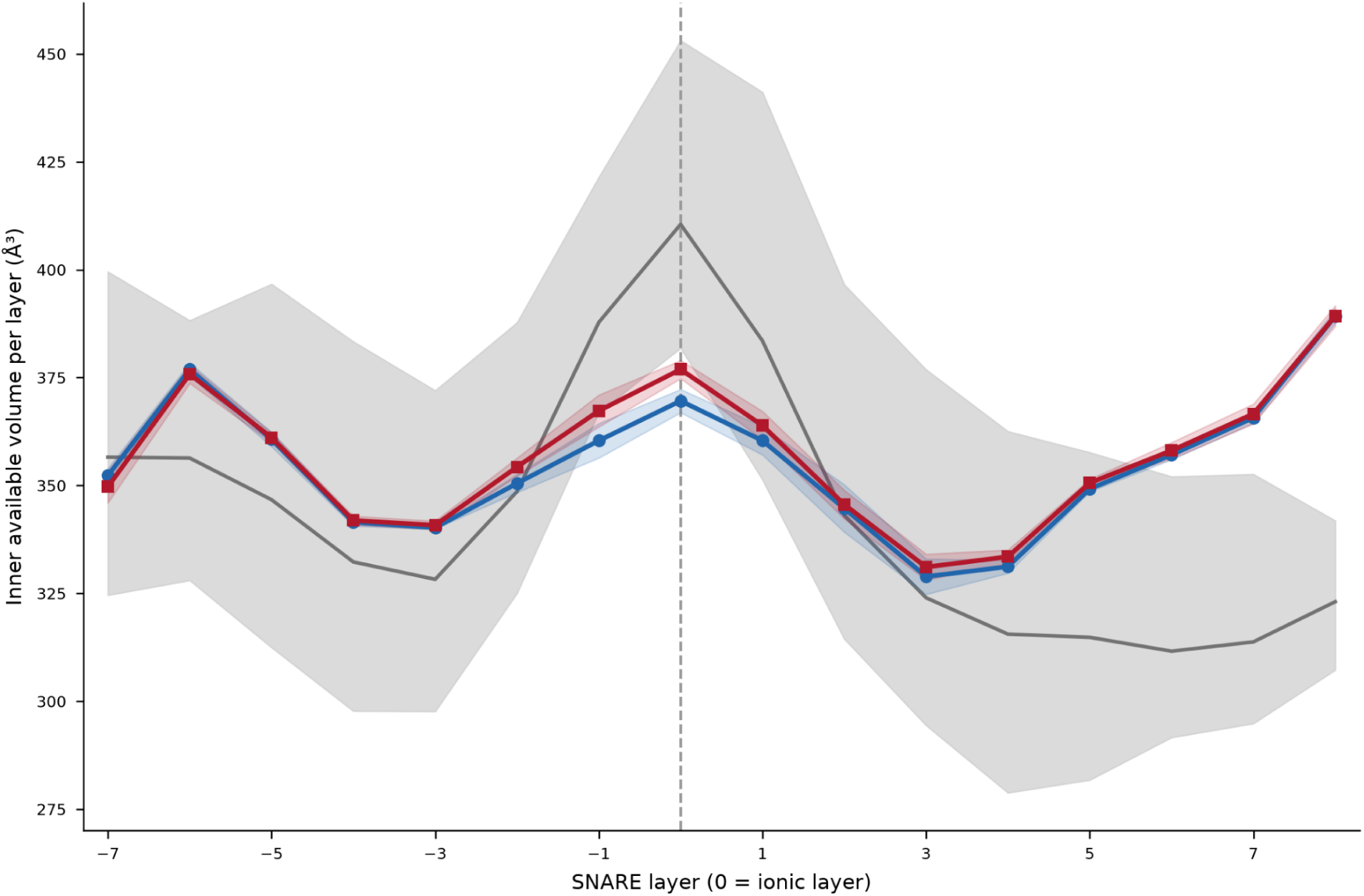
Comparison of the core volumes of the Heimdall and eukaryotic SNARE complexes. The volume occupied by amino acids within the SNARE bundle was calculated for each layer of the four-helix bundle using the structure of the Heimdall SNARE complex (blue), a mutant complex in which one of the four 0-layer glutamines was replaced by arginine (red), and published structures of eukaryotic SNARE complexes (grey). Grey zone, range (min-max) of inner available volume across five eukaryotic SNARE complexes. Across eukaryotic SNARE complexes, the occupied volume varies along the bundle in a characteristic spindle-shaped profile. The Heimdall SNARE complex follows a similar profile, with the occupied volumes at the −1 and 0 layers being slightly lower than those observed in eukaryotic SNARE complexes, while the remaining layers fall within the range observed for eukaryotic complexes.

